# UV-irradiated rotifers for the maintenance of gnotobiotic zebrafish larvae

**DOI:** 10.1101/2024.08.15.608160

**Authors:** Susana Márquez Rosales, Peter I. Bouchard, Emily M. Olmstead, Raghuveer Parthasarathy

## Abstract

Host-associated microbial communities profoundly impact the health of humans and other animals. Zebrafish have proven to be a useful model for uncovering mechanisms of host-microbe interactions, but the difficulty of maintaining germ-free or gnotobiotic zebrafish beyond one week post-fertilization has limited their utility. To address this, we have developed a simple protocol using ultraviolet (UV) irradiation of rotifers, a common and nutrient-rich prey species for larval zebrafish, to reduce the bacterial load associated with the rotifers by several orders of magnitude while maintaining their motility and viability. We find that though feeding with UV-treated rotifers does not preserve the sterility of germ-free fish, it enables the maintenance of pre-existing bacterial communities. Normal feeding, in striking contrast, leads to the near total depletion of these prior populations. We measure the abundance of single- and three-species consortia of zebrafish-commensal bacteria inoculated into initially germ-free larvae in a series of experiments extending to 8 days of feeding, or 13 days post-fertilization. We find, in fish fed UV-treated rotifers, persistence of bacterial populations on timescales of days, together with strong species-specific variation. In addition, re-inoculation of differently labeled strains of the same zebrafish-commensal species alongside feeding leads to colonization by the new bacteria without displacement of earlier microbes. Our method will facilitate the use of gnotobiotic zebrafish for investigations of phenomena that emerge later in animal development and for studies that probe microbiome composition fluctuations and stability over extended timescales.

## Introduction

The human body sustains nearly 100 trillion microorganisms, most of which form dense communities in the intestinal tract, communicating with host organs and influencing the course of health and disease [1]. Manipulating and controlling microbial variables to uncover mechanisms of host-microbe interactions is challenging in humans, which has long motivated the use of animal models and the development of strategies to raise them germ-free, i.e. devoid of microorganisms, or under gnotobiotic conditions, i.e. with known microbial strains or communities. Reports of germ-free guinea pigs date from 1896 [2], and since then, researchers have successfully derived various germ-free animals, including mammals, birds [3], amphibians [4], and fish. In 1942, Baker et al. reported the first germ-free fish (platyfish, *Xiphophorus maculatus*) [5], which was followed by other aquatic species including zebrafish (*Danio rerio*) [6, 7], a species whose attributes such as rapid development, genetic tractability, and transparency at young ages have made it a widely used animal model [8–10]. Zebrafish-based studies of host-microbe interactions have illuminated the varied roles of commensal microbes in early development [9] including effects on epithelium maturation, secretory cell proliferation, enzymatic activity [6, 11–13], immune cell numbers [14–16], the abundance of insulin-producing *β* cells [17, 18], and neural anatomy influencing behavior [19]. Cell-resolved live imaging of gut microbes in larval zebrafish [20], so far impossible in other vertebrates, together with a library of fluorescently-labeled zebrafish-native bacterial species [21], has revealed diverse aggregation states among gut microbes [22–24], influences of bacterial motility on host immune response [25], effects of antibiotics on bacterial motility and persistence [26], and non-pairwise interactions among gut bacterial species [27].

One major aspect of microbiome research to which zebrafish have contributed far less as a model is the investigation of long timescales, necessary for studying bacterial community stability and changes over developmental periods extending to the whole lifespan. This is important because studies in humans show significant variations of timescales bacterial of persistence throughout a host’s life, ranging from mere days or weeks, particularly during early developmental stages [28], to extended periods stretching up to months or even years in adults [29–31]. Perturbations like changes in diet or antibiotic introduction can also alter the microbiota on timescales of days or weeks, though a full return to the original state can take years [32–35]. For all these temporal changes, the underlying mechanisms are poorly understood [36]. Zebrafish could help illuminate these issues for the reasons noted above, but they have suffered from a key limitation: the difficulty of maintaining gnotobiotic animals beyond approximately 7 days post-fertilization (dpf), at which point, having exhausted their yolk, they require food. While it is possible to provide sterilized, powdered chow [6], zebrafish larvae raised on dry food show diminished growth and development compared to larvae fed live food [37]. Larval zebrafish are primarily visual hunters, and the motion of prey is crucial to stimulating capture and food intake [38, 39]. Several research groups have therefore pursued the derivation of germ-free prey species. These include *Artemia* (brine shrimp), which researchers since the 1950s have made germ-free through egg sterilization [40–43]. Recently, Jia *et al*. [44] showed that a different fish species (medaka, *Oryzias melastigma*) could be raised germ-free to adulthood solely on *Artemia*. Zebrafish are capable of eating *Artemia* by 9 or 10 dpf, a few days after yolk depletion, calling for another, intermediate, prey species. For earlier feeding of zebrafish, groups have reported antibiotic treatment of the unconventional prey species *Tetrahymena thermophila* [41,42]. While successful for producing gnotobiotic fish, such methods involving these prey species, especially *Artemia*, are highly labor intensive [7] and are not widely used. Moreover, antibiotics, even at low concentrations, can have large impacts on the gut microbiome [26, 35] and can alter animal metabolism and growth [45], further motivating our efforts to investigate alternative methods. We consider rotifers (*Brachionus plicatilis*), an easily grown and widely used live food for zebrafish larvae [46, 47], and ultraviolet (UV) irradiation. Rotifers also have been the target of axenic method development for over half a century at least [48], for example through sterilization of amictic eggs [49], though typically the aim is to facilitate studies of rotifers themselves rather than their use as prey. Ultraviolet sterilization of rotifers has rarely been investigated.

In 1999, Munro et al. reported an ultraviolet irradiation protocol that reduced the bacterial load of rotifers by 99% and showed that these rotifers could be successfully fed to turbot [50]. No investigations have been made, however, on the gnotobiotic status of fish fed UV-irradiated food. We therefore aimed to establish a simple method based on UV-exposed rotifers that can maintain gnotobiotic zebrafish for extended durations. We developed and compared two separate procedures to minimize native bacterial populations in rotifers: antibiotic treatment and a simple UV irradiation protocol. We assessed the effect of consumption of treated rotifers on pre-existing bacterial populations in larval zebrafish. While both prey treatments reduced the rotifers’ bacterial load, consumption of antibiotic-treated rotifers led to dramatic drops in commensal bacterial abundance. In contrast, consumption of UV-treated rotifers left larval bacterial populations largely intact. Then, we examined bacterial populations in fed larvae to 13 dpf, demonstrating greater stability in fish fed UV-treated rotifers than in untreated rotifers. The methods described here will facilitate studies of gut microbiome dynamics and host-microbe interactions in gnotobiotic animals.

## Results

We aimed to establish an easily implemented method for feeding rotifers to zebrafish larvae that maintains gnotobiotic conditions. The key step is to reduce as much as possible the bacteria that naturally accompany the rotifers while preserving rotifer survival and motility. We investigated two methods: one using ultraviolet light exposure and the other using antibiotics.

The ultraviolet irradiation method uses a 270-280 nm UV-C LED attached to the cap of a flask containing a rotifer suspension and a stir bar to ensure mixing (Figure 1A; see Materials and Methods). To find an exposure protocol that leads to a significant reduction in bacterial numbers while maintaining a viable rotifer population, we assessed cycles of UV light consisting of alternating 30-minute exposure and rest periods. By three exposure cycles, the plating of the rotifer suspension shows a drop in bacterial abundance of nearly four orders of magnitude (Figure 1B).

**Figure 1.**
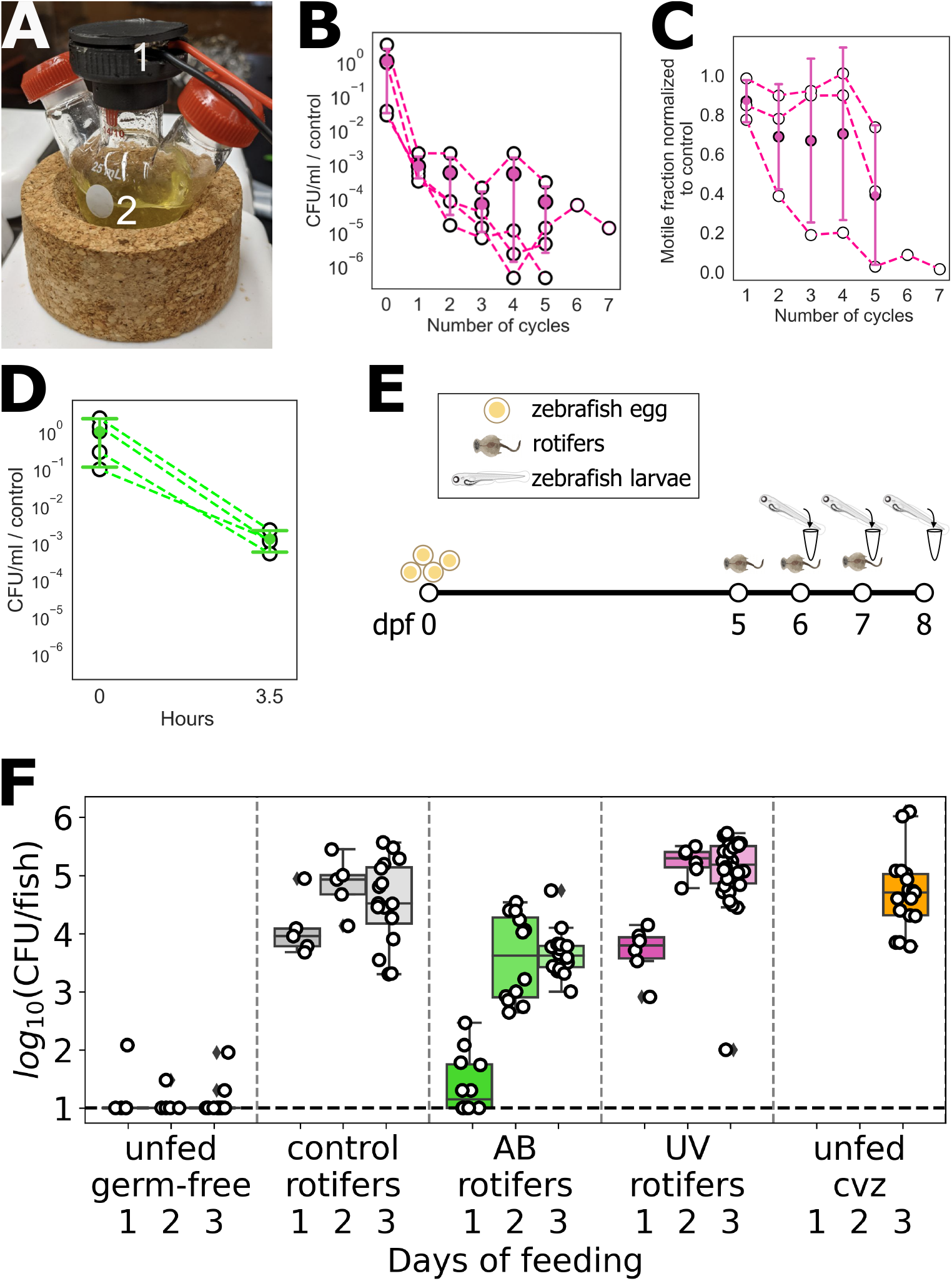
**A** Experimental setup for UV irradiation of rotifers, including (1) a UV-C LED mounted in a cap of a round bottom flask and (2) rotifers in a 4ppt NaCl solution. **B** Bacterial concentration in the rotifer suspension as a function of the number of UV exposure cycles, each 30 minutes on / 30 minutes off. **C** Fraction of motile rotifers as a function of the number of UV exposure cycles. In B and C, open symbols represent different replicates, and solid symbols and error bars indicate the mean and standard deviation. **D** Bacterial concentration in the rotifer suspension before and after incubation with antibiotics for 3.5 hours. **E** Schematic diagram of the protocol for assessing bacterial load in initially germ-free zebrafish subject to different feeding methods. **F** Bacterial abundance per larval zebrafish after 1, 2, and 3 days of feeding with rotifers under different treatments, or unfed at the same days. Each symbol indicates a measurement from an individual fish; boxes indicate the median (line within the box), first and third quartiles (top and bottom of the box), and 95% confidence intervals (bars). The dashed line indicates the approximate limit of detection, 10 bacteria per fish.

UV irradiation causes rotifers to become slow or non-motile, with the non-motile ones possibly being dead. By five UV cycles, the fraction of motile micro-animals decreases to less than 50%, and the average speed of motile rotifers falls to less than half that of unexposed rotifers (Fig. 1C, Supplemental Fig. 1, Supplemental Movies S1, S2; see Materials and Methods). We therefore chose four UV exposure cycles to reduce the rotifers’ bacterial load while keeping the majority of them motile.

While a large variety of antibiotics exist, we focus for simplicity on the same cocktail of antibiotics typically used for germ-free zebrafish derivation [7] (see Materials and Methods). After 3.5 hours of incubation, the antibiotics reduce the bacterial load of rotifer suspensions by three orders of magnitude (Fig. 1D), similar to the results from UV irradiation. Following antibiotic treatment, we observed no impact on rotifer swimming speed or the proportion of motile rotifers.

We evaluated the efficacy of the UV and antibiotic treatments for maintaining germ-free fish after feeding. Beginning with germ-free larvae, we administered feeding regimes spanning one, two, and three days starting at 5 dpf (Figure 1E), followed by bacterial abundance quantification via larval homogenization and plating (see Materials and Methods). Notably, we measured bacterial load from entire larvae, which will be dominated by gut microbes but can include skin-resident microbes. Figure 1F presents the results detailing bacterial loads from larvae fed with UV-treated rotifers, antibiotics-treated rotifers (AB), and control (untreated) rotifers, alongside unfed germfree fish and unfed conventionalized (CVZ) larvae, the latter initially germ-free and then raised in standard, non-sterile aquaculture water. As expected, unfed conventionalized larvae and larvae fed control rotifers show 10^4^-10^5^ bacteria (colony forming units (CFU)) per fish, while unfed germ-free fish typically have undetectable numbers of bacteria (Figure 1F). Fish fed with AB-treated rotifers showed low bacterial abundance after one day of feeding (10-100 CFU), though bacterial numbers increased to several thousand by two or three days of feeding. Fish fed with UV-treated rotifers lost their germ-free status after just one day of feeding, with bacterial abundance around 10^4^ indicating that the small amount of bacteria remaining in the rotifers after UV treatment are sufficient to colonize the larvae, compromising the gnotobiotic conditions. We conclude that maintaining germfree fish after feeding is not feasible with these approaches. We also measured larval length after two or three days of feeding as an indicator of fish health. Larvae in all fed groups were, on average, longer than unfed larvae (Supplemental Figure 2). The length of fish fed with UV-treated rotifers showed no significant difference compared to those fed with control rotifers, implying that the decrease in motility and speed of rotifers due to UV treatment did not negatively affect their capture by hunting fish.

**Figure 2.**
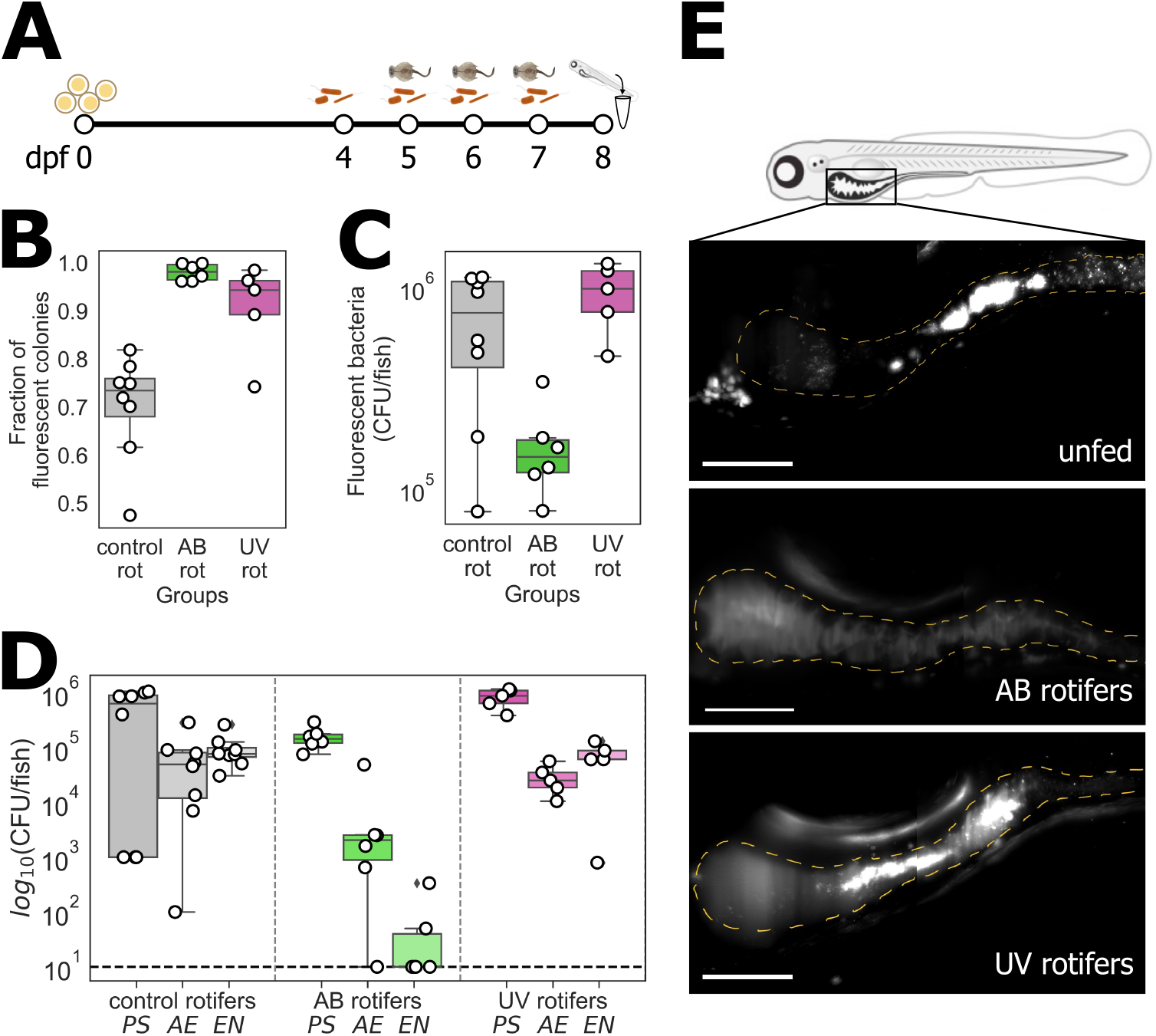
**A** Schematic diagram of the protocol to evaluate the permanence of gnotobiotic larvae after three days of feeding, in which initially germ-free zebrafish are inoculated with three fluorescently labeled bacterial species. **B** Fraction of fluorescent colonies from plating whole larvae, indicating the fraction of intentionally inoculated bacteria, after three days of feeding and re-inoculation. **C** Total abundance of fluorescent bacteria per larva after three days of feeding and re-inoculation. **D** Abundance of each intentionally inoculated strain. PS: *Pseudomonas mendocina* ZWU0006, AE: *Aeromona veronii* ZOR0001, EN: *Enterobacter cloacae* ZOR0014. In B-D, each symbol indicates a measurement from an individual fish; boxes indicate the median and quartiles. **E** Maximum intensity projections of representative 3D image stacks from light sheet fluorescence microscopy of the guts of live larval fish, taken after one day of feeding with antibioticor UV-treated rotifers, or on the equivalent day for unfed fish. Dashed lines trace approximate intestinal boundaries. Bar = 200 μm.

Though the methods being examined cannot maintain zebrafish germ-free, they nonetheless may be capable of keeping larvae under gnotobiotic conditions, sustaining intentionally inoculated bacterial communities with minimal perturbation. We hypothesized that symbiotic strains may out-compete the bacteria that accompany rotifers. To assess this, as indicated schematically in Figure 2A, we started by establishing a bacterial community at 4 dpf by inoculating germ-free larval fish with previously studied zebrafish-native strains expressing fluorescent proteins (see Materials and Methods) [21, 27] *Pseudomonas mendocina* ZWU0006 (PS), *Aeromona veronii* ZOR0001 (AE), and *Enterobacter cloacae* ZOR0014 (EN). On each of the next three days (5 dpf), we fed the fish and also re-introduced the fluorescent strains to the aqueous medium. At dpf, we homogenized and plated the whole fish as above (see Materials and Methods), noting the abundance of fluorescent colonies, representing the intentionally inoculated species, and nonfluorescent colonies, representing the species introduced by feeding. We found that for fish fed AB-treated and UV-treated rotifers, the fraction of fluorescent bacteria is over 90% (0.98 ± 0.016, mean ± standard deviation) and 0.90 ± 0.087, respectively) (Figure 2B). In contrast, the fraction of fluorescent bacteria in fish fed control rotifers is considerably smaller, 0.7 ± 0.1. Strikingly, the overall number of intentionally inoculated (fluorescent) bacteria was approximately an order of magnitude lower in fish fed AB-treated rotifers than in fish fed UV-treated or control rotifers (Figure 2C). Plating on chromogenic agar, on which colonies of the different intentionally inoculated strains show different colors [27] (see Materials and Methods) shows that the depletion of bacterial numbers following feeding with AB-treated rotifers is strongly species dependent (Figure 2D), with the EN population being particularly affected and completely absent in most larvae (Figure 2D). The AE population is two orders of magnitude smaller compared to other fed groups, while PS is similar (Figure 2D). To gain insights into intestinal abundance in particular, we imaged the whole intestines of live larvae using Light Sheet Fluorescence Microscopy [22, 24]. We could not detect fluorescent bacteria in the gut of larvae fed with AB-treated rotifers, while bacteria were clearly evident in larvae fed with UV-treated or control rotifers (Figure 2E). Our observations indicate the feasibility of keeping fish under gnotobiotics conditions with UV-treated rotifers, which largely preserve zebrafish-associated bacterial communities.

Having established a method using UV irradiated rotifers to feed zebrafish larvae, we investigated inoculated *Enterobacter cloacae* (EN) populations over several days, noting that of the three strains examined this strain is the most sensitive to food-induced perturbation (Fig. 2D). To distinguish bacteria descended from those inoculated on the first day, colonizing germ-free fish, from bacteria introduced on subsequent days, we used differently tagged EN strains [21], expressing green fluorescent protein (GFP) and dTomato, respectively. As above, non-fluorescent bacteria represent unintended colonizers from rotifers or other sources. Fish were fed from 5 to 7 dpf or from 7 to 9 dpf. The first set allows us to include unfed fish as a control group. The second set tests bacterial dynamics into the regime that requires feeding. For both sets, we include fish fed with control (untreated) rotifers to determine the efficacy of the UV treatment.

The study comprises four distinct fish groups for each set of days, as illustrated in Figure 3A. All fish were inoculated on the initial day (4 or 6 dpf) with GFP-labeled EN, fed with UV-treated or control rotifers on each of the next three days unless in the unfed set, and then plated to determine bacterial load on various days including the final day (Fig. 3A). Groups 2, 3, and 4 were, in addition, inoculated with dTomato-labeled EN on the first, second, and third day of feeding, respectively (Fig. 3A; see Materials and Methods). Representative larvae were homogenized and plated as above to quantify bacterial load, with all fish assessed by the final day, at 8 or 10 dpf. Together, these groups provide insight into the persistence of *Enterobacter* introduced before feeding and of stability following re-inoculation.

**Figure 3.**
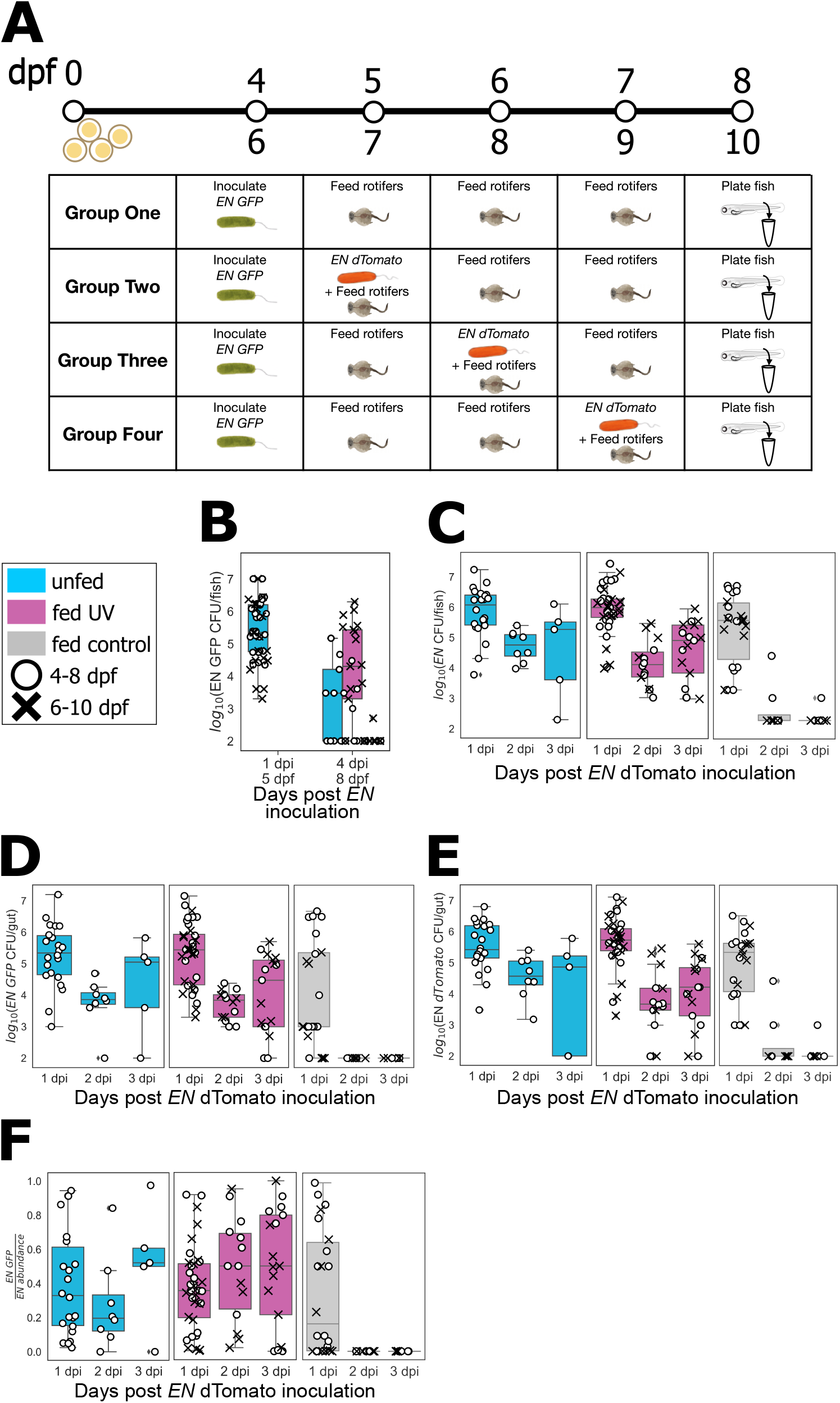
**A** Schematic diagram of the protocol to evaluate *Enterobacter* stability after feeding and re-inoculation. As described in the main text, initially germ-free fish were inoculated with GFP-labeled EN and fed UV-treated or control rotifers or left unfed for three days, starting at 4 or 6 dpf. Groups 2-4 were re-inoculated with dTomato-labeled EN on different days. **B F** Final EN abundance in unfed larvae (cyan), larvae fed with UV treated rotifers (pink), and larvae fed with untreated rotifers (gray). Circles represent data from larvae fed from 5 to 7 dpf, and crosses from 7 to 9 dpf. **B** Fluorescent EN-GFP abundance in Group 1 larvae assessed one day post-inoculation (before the start of feeding) and four days post-inoculation (after three days of feeding). **C**-**E** From groups 2, 3, and 4, total fluorescent EN abundance (GFP and dTomato) (C), EN-GFP abundance (D), and EN-dTomato abundance (E) one, two, or three days post EN dTomato inoculation. F Fraction of EN-GFP relative to the total EN abundance one, two or three days post EN-dTomato inoculation.

We present the results in Figure 3, indicating the feeding types by different colors and the sets of days by different symbols. Considering fish only inoculated on the first day, by EN-GFP, the initial bacterial load assessed from unfed fish at one day post inoculation (dpi) was roughly 10^5^-10^6^ per fish, as expected (Fig. 3B). At 4 dpi, almost no *Enterobacter* were found in fish fed with untreated rotifers, in contrast to roughly 10^3^-10^5^ in fish fed UV-treated rotifers. Interestingly, unfed fish at 8dpf showed fewer *Enterobacter* (Fig. 3B), suggesting that the depletion of yolk has consequences for resident bacteria. We next consider the total EN population in Groups 2, 3, and 4, as a function of days following inoculation of the second (EN-dTomato) bacterial strain (Fig. 3C). Again, almost no *Enterobacter* were found in fish fed with untreated rotifers at 2 and 3 dpi, while unfed fish and fish fed UV-treated rotifers showed sizable bacterial numbers (10^4^-10^5^ per fish, Fig. 3C). Considering separately the EN-GFP population, descended from the initial colonizers, and the EN-dTomato population, descended from later entrants, we find similar orders-of-magnitude for the abundance of each group in the unfed fish and fish fed UV-treated rotifers (Fig. 3D, E). The relative abundance of the two groups, quantified as the GFP-labeled fraction of the total *Enterobacter* abundance, shows considerable variation (Fig. 3F), indicating that neither the initial nor later colonizers consistently dominates the population.

Overall, the similarity between the abundances of intentionally inoculated bacteria in unfed fish and those fed with UV-treated rotifers, and the marked contrast with fish fed untreated rotifers, validates the UV treatment protocol for studying zebrafish-associated bacterial stability.

Finally, we examined the stability of a multi-species bacterial consortium in zebrafish up to 13 dpf, several days longer than is possible without feeding. We again consider PS, AE, and EN (as in Fig. 2) and again consider differently inoculated groups. We inoculated all larvae with GFP-labeled PS, AE, and EN at 4 dpf and Group 2 larvae additionally with dTomato-labeled PS, AE, and EN at 8 dpf, the fourth day of feeding (Fig. 4A; see Materials and Methods). Both groups were fed for 7 or 8 days after which individuals were homogenized and plated to assess bacterial load. Both groups included fish fed UV-treated rotifers and control (untreated) rotifers.

**Figure 4.**
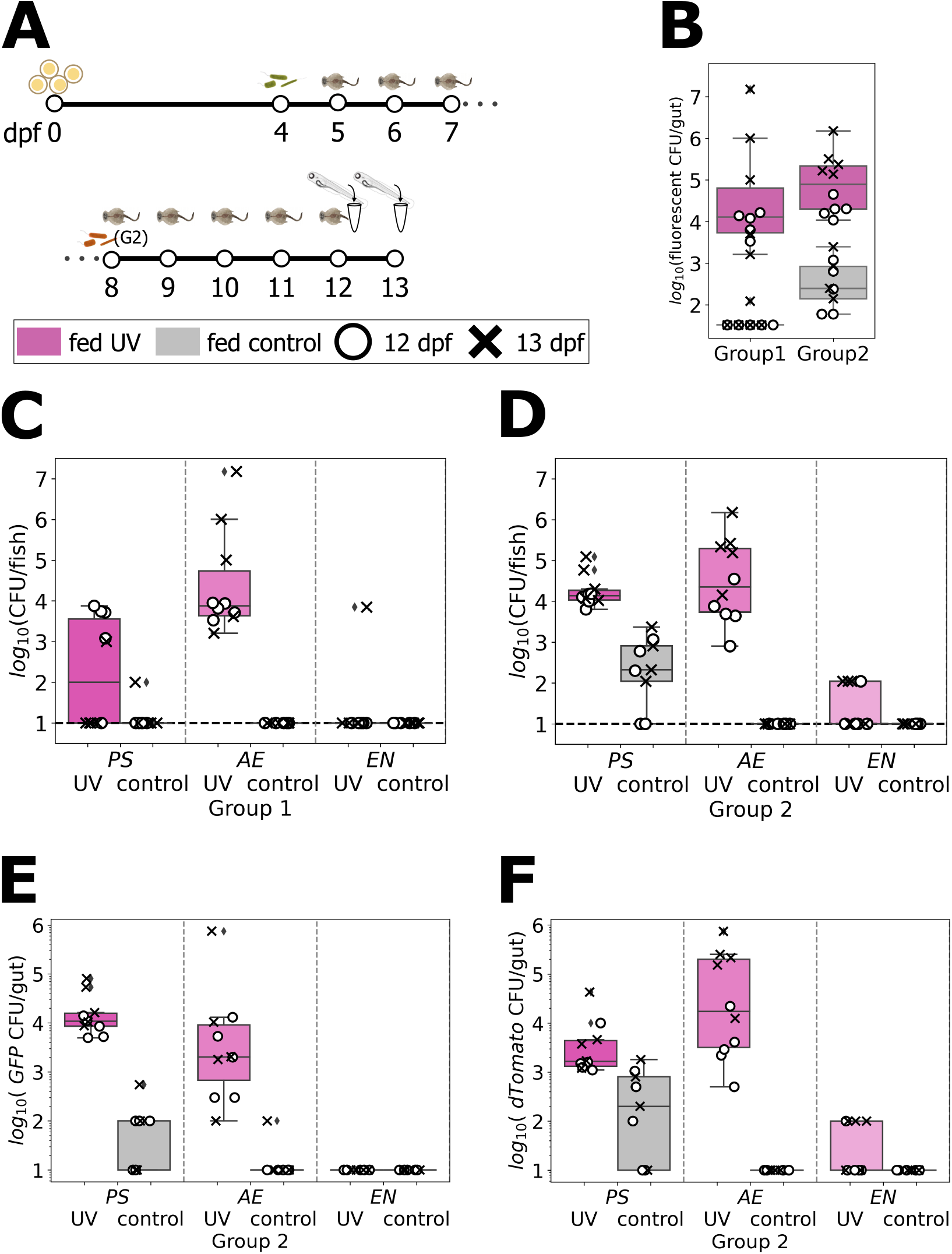
**A** Schematic diagram of the protocol to evaluate the stability of multiple bacterial species in larvae fed until 13 dpf. Initially germ-free larvae were inoculated with GFP-labeled PS, AE, and EN at 4 dpf and split into two groups. Zebrafish in Group 2 were re-inoculated with dTomato-labeled PS, AE, and EN at 8 dpf. Both groups were fed from 5 to 11 or 12 dpf and homogenized and plated at 12 or 13 dpf. **B** Fluorescent bacterial abundance per larva after 7 or 8 days of feeding. Pink corresponds to larvae fed with UV treated rotifers, and gray to larvae fed with untreated rotifers. **C** and **D** Bacterial abundance of each inoculated per strain for fish in Group 1 (C) and Group 2 (D). **E** Total abundance of GFP-labeled bacteria, descended from initial colonizers, of each strain in larvae from Group 2. **F** Total abundance of dTomato-labeled bacteria, descended from later colonizers, in larvae from Group 2.

Considering the total abundance of fluorescent (intentionally inoculated) bacteria at the experimental endpoint, fish in Group 1 and Group 2 that were fed UV-treated rotifers showed sizeable populations, roughly 10^4^-10^5^ CFU/fish, while fish fed with control rotifers showed bacterial numbers that were on average two orders of magnitude lower (Fig. 4B). Decomposing this by inoculated bacterial species shows considerable differences (Fig. 4 C, D). AE was the most stable strain, with abundances similar to those measured one day after bacterial inoculation in earlier experiments (Fig 2D), and EN was the least stable strain. For all strains, abundances in control-rotifer-fed fish were low or undetectable (Fig. 4 C, D). Considering separately, in Group 2 fish, the GFP-labeled population, descended from the initial colonizers, and the dTomato-labeled population, descended from later entrants, we find similar orders-of-magnitude for the abundance of each in the fish fed UV-treated rotifers (Fig. 4D, E). Intriguingly, PS and AE show opposite behaviors for the abundance of dTomato-relative to GFP-labeled bacteria, with PS dominated by the descendants of initial colonizers and AE the later colonizers (Fig. 4 E, F). Variation is large, however, prohibiting strong conclusions.

## Discussion

To improve the utility of zebrafish as a model for studying animal associated microbiomes, we have developed a method for feeding zebrafish larvae that allows live, motile food while minimizing the disruption of existing bacterial communities. Our method uses UV-C LEDs that are inexpensive, easy to deploy, and sufficient to reduce within a few hours the bacterial density that naturally accompanies rotifers by four orders of magnitude. Although maintaining truly germ-free zebrafish larvae is not achievable with this method of preparing prey, likely due to the ability of remaining microbes to proliferate in the sterile larval environment, we find that feeding with UV-irradiated rotifers allows zebrafish-native bacteria associated with the fish prior to feeding, or reintroduced together with feeding, to persist at high abundance. Normal feeding on untreated rotifers, in contrast, diminishes fish native bacteria by orders of magnitude.

Treatment of rotifers with an antibiotic mixture commonly used for deriving germ-free fish, though effective for reducing the bacterial concentration in the rotifer suspension (Fig. 1D) leads after ingestion to large and species-specific drops in fish-associated bacterial abundance (Fig. 2 C, D). This is perhaps unsurprising, as the antibiotics are diluted only by a factor of 15 from the concentration used for deriving germ-free fish (see Materials and Methods). It may be possible, for example through filtration or centrifugation, to separate the rotifers from their antibiotic treated suspension prior to feeding, but this adds extra procedural steps, and moreover it is possible that the antibiotics are preferentially concentrated in the rotifers. Furthermore, studies of humans [51] and zebrafish [26] have shown that even low doses of antibiotics can have large consequences on gut microbiomes, in part due to antibiotic-induced changes in bacterial morphology and motility [26], providing further cautions for antibiotic-based procedures for gnotobiotic prey. We believe, therefore, that our use of UV-irradiation highlights a gnotobiotic strategy that is less likely to lead to strong perturbations of animal-associated microbiomes, and we focus on further characterization of this method.

Extending the timescale over which zebrafish can be maintained under gnotobiotic conditions opens up new research opportunities. For example, week-old gnotobiotic zebrafish larvae have enabled insights into connections between gut bacterial species and innate immune responses (e.g. [52]), but studying the adaptive immune system requires older fish, as the maturation of the adaptive immune system in zebrafish starts at three weeks post-fertilization with the appearance of CD4+/CD8+ lymphocytes [53]. Given increasingly appreciated connections between hostassociated microbes and autoimmune and chronic inflammatory disorders [54], older gnotobiotic zebrafish could serve as valuable experimental tools.

Most generally, improved gnotobiotic methods enable the extension of timescales of experimental investigation in zebrafish. Of particular interest are timescales of gut microbiome stability and instability, whose underlying determinants in humans and other animals remain mysterious [36]. Given the presence of a bacterial strain, what is the likelihood of its persistence some time later, and is this altered by immigration of members of the same or different strains?

We have demonstrated that studying these questions is tractable, for example examining a three-species consortium in larvae fed for up to eight days, to 13 dpf. We find strong species-dependence of stability, with a zebrafish-native *Enterobacter* nearly disappearing while other species persist (Fig. 4). Notably, this *Enterobacter* species also shows the greatest decline when fish are fed with antibiotic-treated rotifers (Fig. 2D). These observations are consistent with previous results from short-duration studies that show that *Enterobacter* is disfavored in competition with *Aeromonas* [23, 27] and is highly sensitive to weak antibiotics [26]. *Enterobacter* is also highly aggregated [22], and we suspect that underlying all of these phenomena is a connection between aggregation and intestinal transport. With the ingestion of food, a frequent perturbation of obvious importance to humans and other animals, the physical space occupied by food particles and the mechanics of its digestion may induce species-specific displacement, particularly impacting more aggregated strains. Further experiments, including studies using live imaging (as in Fig. 2E), can directly probe and quantify these processes in a useful model animal.

## Materials and Methods

### Animal care

All experiments with zebrafish were done in accordance with protocols approved by the University of Oregon Institutional Animal Care and Use Committee and by following standard protocols [55].

### Zebrafish gnotobiology

Wild-type ABC X TU zebrafish (*Danio rerio*) were derived germ-free as described previously [7] with slight modifications. In brief, fertilized eggs were collected and placed in sterile antibiotic embryo medium (EM) containing 100 μg/mL ampicillin, 250 ng/mL amphotericin B, 10 μg/mL gentamicin, 1 μg/mL tetracycline, and 1 μg/mL chloramphenicol for approximately 5 hours. The eggs were then washed in sterile EM containing 0.003 % sodium hypochlorite and then in sterile EM containing 0.1 % polyvinylpyrrolidone-iodine. Washed embryos were distributed into tissue culture flasks containing 50 mL of sterile embryo medium at a density of one embryo per mL. Flasks were inspected for sterility before being used in experiments.

### Bacterial strains

We used the following strains, each originally isolated from the zebrafish intestine: *Aeromonas* sp. strain ZOR0001, *Pseudomonas mendocina* (ZWU0006), and *Enterobacter* sp. strain ZOR0014 (EN). From each, engineered strains constitutively expressing green fluorescent protein (GFP) or dTomato were previously generated ([21]). Stocks of bacteria were maintained in 25% glycerol at ≥80 ^°^C.

### Inoculation of larval zebrafish

One day prior to fish inoculation, bacteria from frozen glycerol stocks were grown overnight in lysogeny broth (LB medium; 10 g/L NaCl, 5 g/L yeast extract, 12 g/L tryptone, 1 g/L glucose) at 30 ^°^C with shaking. For inoculation of germ-free fish with a single species (*Enterobacter* sp. strain ZOR0014), 1 mL of overnight culture was washed once by centrifuging for 2 minutes at 7000 x g, removing the supernatant, and adding 1 mL of fresh sterile embryo medium. Then 100 μL of the bacterial mix was added to a 50 mL tissue culture flask containing approximately 50 germ-free 4 dpf zebrafish larvae, giving a bacterial concentration of approximately 10^6^ CFU/mL. For experiments involving multiple bacterial species, we combined 1 mL of each overnight culture, diluting each as needed in lysogeny broth to achieve similar optical densities (OD600 ≈ 5.0) and therefore similar concentrations. The bacterial mixture was further prepared and added to the flasks containing larvae as above, at a bacterial concentration of approximately 10^6^ CFU/mL. For re-introduction of bacteria to previously colonized fish, we prepared a 1000-fold dilution of the *Enterobacter* overnight culture in LB for single-species experiments and a 100-fold dilution of the bacterial mixture for multi-species experiments. Then, 100 μL of the bacterial suspension was combined with 900 μL of the rotifer suspension and added to the flasks containing zebrafish larvae as above, giving bacterial concentrations of approximately 10^4^ CFU/mL and 10^5^ CFU/mL (about 10^4^ CFU/mL per each species) for the single- and multi-species experiments, respectively.

### Rotifers

Rotifers (*Brachionis plicatilis*) were provided by the University of Oregon Zebrafish Facility. Rotifer cultures were raised in the facility in 5 gallon containers, fed with the “Rotigrow Plus” algae mixture (Reed Mariculture), and maintained at 4 ppt salinity in “Instant Ocean” commercial sea salt mix (Instant Ocean). Rotifers at a density of roughly 2000/mL were obtained as needed from the facility and used the same day.

### Ultraviolet irradiation of rotifers

Ultraviolet light, 270-280 nm, was provided by LEDs (E275-3-S UVC LED Chip on Board, International Light Technologies, $16.85 each; UVC LED Driver ILT-PWRTYLED.3W, International Light Technologies, $21.91). An LED was glued to a 3D-printed cap designed to fit a three-neck round bottom flask (Synthware Round Bottom Flasks, Three Neck, Threaded, 25 mL; Fisher Scientific Catalog No.31-502-121); CAD files for the cap are provided as Supplemental Material. The remaining necks of the flask, which allow easy extraction of the rotifer suspension, were kept loosely capped to allow oxygenation. For rotifer treatment, rotifers were diluted 5X in 4 ppt NaCl, reducing the opacity of the suspension, to a concentration of roughly 400/mL and 15 mL of this suspension was placed in each round bottom flask. A stir bar, 2 mm, was used for continuous stirring. The UV intensity was measured to be 0.48 mW/cm^2^ at the location of the water surface, 5.4 cm away from the LED.

### Antibiotic treatment of rotifers

We use the same antibiotic solution and concentrations noted in the Zebrafish Gnotobiology section, treating 1 mL of rotifer suspension for 3.5 hours.

### Rotifer motility assessment

Rotifer motility was assessed by tracking individual rotifers in brightfield movies captured at 10 frames per second with a scale of 6.5 μm/px. Using custom software, images were inverted and regions of interest containing rotifers were identified as local intensity maxima. Rotifers were localized in each frame using a symmetry-based algorithm [56] and connected across frames using nearest-neighbor linkage. A rotifer was considered motile if the mean speed of its trajectory exceeded 50 μm/s, which clearly distinguished moving and stationary rotifers.

### Feeding

Before feeding, larvae were transferred to a new tissue culture flask with fresh sterile embryo medium. To 15 mL of the flask medium, we added 1 mL of rotifer suspension, corresponding to roughly 400 rotifers, or roughly 30 per fish. Fish were allowed to feed *ad libitum*.

### Light sheet fluorescence microscopy

Imaging was performed using a home-built light sheet fluorescence microscope based on the design of Keller et al. [57]. The protocols to mount the fish and the details of the microscope can be found in [58], [59], [60], and [25]. In brief, we anesthetize larvae with MS-222 (tricaine, 20 μL/mL) and mount the fish in glass capillaries containing 0.7% agarose gel suspended vertically, head up, in a custom imaging chamber containing EM and MS-222. Lasers with excitation wavelengths of 488 and 568 nm were used to excite green fluorescent protein and dTomato labeled bacteria, respectively. To image the entire extent of the intestine covering the gut (approximately 1200×300×150 μm) we move the specimen in z-steps of 1 μm and sequentially image four or five sub-regions computationally registering the images after acquisition.

### Plating and bacterial abundance measurement

To quantify bacterial abundance in the fish, we euthanize the fish via hypothermic shock and transfer single larva into 1.5 mL Eppendorf tubes containing 1 mL of sterile embryo medium and approximately 0.1 mL of 0.5 mm zirconium oxide pellets. Whole fish were homogenized with a bullet blender (Next Advance) for 3 minutes at speed 8. Dilutions of 10^−1^ and 10^−2^ were prepared, and 100 μL of each were spread on LB agar plates. Colony-forming units (CFUs) on the plates were counted to quantify bacterial abundances. For experiments with zebrafish inoculated with more than two species, homogenized larvae were plated on Universal HiChrome agar (Sigma Aldrich) to distinguish species based on a colorimetric indicator. All plating-derived abundance data is provided as Supplemental Material.

## Supporting information

Plotted datapoints

STL files for cap to mount UV LED

## Acknowledgments

We thank Rose Sockol, Tim Mason, and the University of Oregon Zebrafish Facility staff for fish husbandry, providing rotifers, and general advice. We also thank Jonah Sokoloff for experimental assistance and Brendan Bohannan, Judith Eisen, Karen Guillemin, and John Rawls for advice and feedback.

This work was supported by the New Tools for Advancing Model Systems in Aquatic Symbiosis program of the Gordon and Betty Moore Foundation (doi:10.37807/GBMF9320). Additional support was provided by the National Science Foundation under the Research Traineeship (NRT) Award DGE 2022168. The funders had no role in study design, data collection and analysis, decision to publish, or preparation.

## Supplemental Material

All plotted datapoints are provided in a CSV file as Supplemental Material. In addition, Supplemental Material includes Supplemental Movies 1 and 2, and a PDF containing Supplemental Figures 1 and 2 and the two movie captions.

