## Supplemental Figures and Movie Captions for "UV-irradiated rotifers for the maintenance of gnotobiotic zebrafish larvae"

#### Contents

- Supplemental Movie Captions.
- Supplemental Figures and Captions.

#### Supplemental Movie Caption

**Supplemental Movie 1.** Representative movie of rotifers without UV irradiation. Rotifer motility is evident, as are a large number of dead rotifers. Brightfield, 10 fps.

**Supplemental Movie 2.** Representative movie of rotifers following four cycles of UV irradiation, as described in the text. Rotifer motility at reduced speed compared to control rotifers is evident, as are a large number of dead rotifers. Brightfield, 10 fps.

### Supplemental Figures

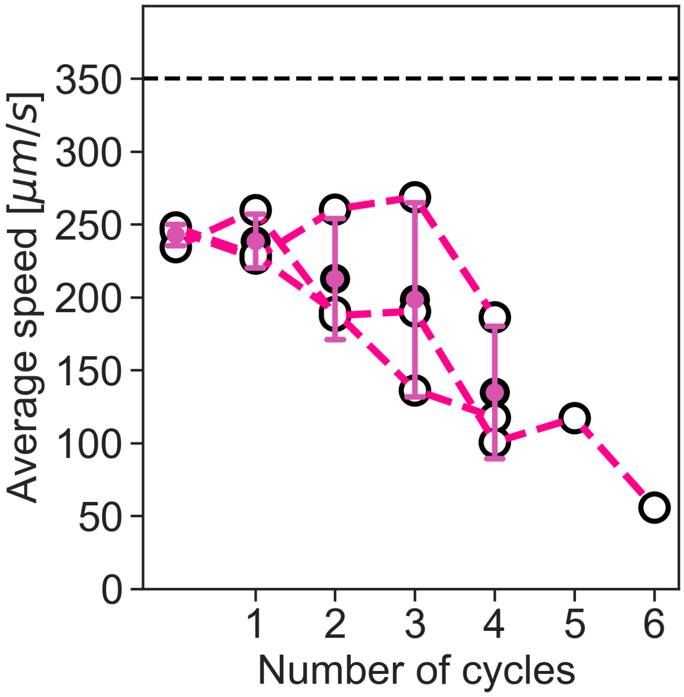


**Supplemental Figure S1**. Average speed of motile rotifers as a function of the number of UV exposure cycles, each 30 minutes on / 30 minutes off. Dotted line at 350μm/s represents the average speed of rotifers before UV irradiation. Open symbols represent different replicates, and solid symbols and error bars indicate the mean and standard deviation.

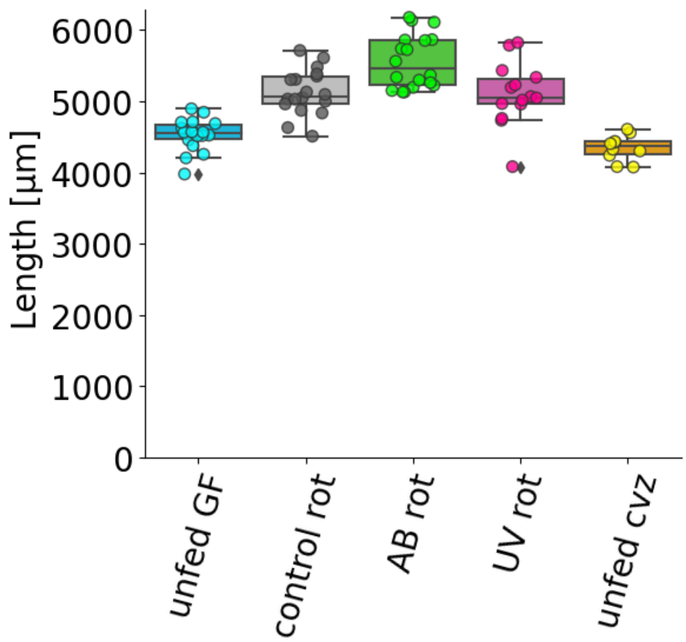


**Supplemental Figure S2.** Fish length on 7 or 8 dpf after 2 or 3 days of feeding with rotifers under different treatments, or unfed at the same days. Each symbol indicates a measurement from an individual fish; boxes indicate the median (line within the box), first and third quartiles (top and bottom of the box), and 95\% confidence intervals.
